# Potential climate change effects on the distribution of urban and sylvatic dengue and yellow fever vectors

**DOI:** 10.1101/2023.11.06.565841

**Authors:** Alisa Aliaga-Samanez, David Romero, Kris Murray, Marina Segura, Raimundo Real, Jesús Olivero

**Affiliations:** Grupo de Biogeografía, Diversidad y Conservación, Departamento de Biología Animal, Universidad de Málaga, Facultad de Ciencias, 29071 Malaga, Spain; Medical Research Council Unit the Gambia at London School of Hygiene and Tropical Medicine, Fajara, Gambia; Centre on Climate Change and Planetary Health, London School of Hygiene & Tropical Medicine, London, UK; Centro de Vacunación Internacional, Ministerio de Sanidad, Consumo y Bienestar Social, Estación Marítima, Recinto del Puerto, Muelle 3, 29001 Malaga, Spain; Instituto IBYDA, Centro de Experimentación Grice-Hutchinson, Malaga, Spain

**Keywords:** Mosquito species, Spatial distribution, Vector-borne diseases, Zoonoses

## Abstract

Climate change may increase the risk of dengue and yellow fever transmission by urban and sylvatic mosquito vectors. Previous research primarily focused on *Aedes aegypti* and *Aedes albopictus*. However, these diseases involve a complex transmission cycle in which sylvatic vectors are also involved. Our aim was to analyse which mosquito species could contribute to the increased risk of transmission of these diseases due to climate change, and to identify where the risk increase could most likely occur. Using a biogeographical approach, we mapped areas where mosquito favourability could increase, decrease or remain stable in the near (2041-2060) and distant (2061-2080) future.

Models predict dengue vectors expanding in West and Central Africa and in South-East Asia, reaching Borneo. Yellow fever vectors could spread in West and Central Africa and in the Amazon. In Europe, the models suggest a re-establishment of *Ae. aegypti*, while *Ae. albopictus* will continue to find new favourable areas. The results underline the need to focus more on vectors *Ae. vittatus*, *Ae. luteocephalus* and *Ae. africanus* in West and Central sub-Saharan Africa, especially Cameroon, Central Africa Republic, and northern Democratic Republic of Congo; and suggest the need for a protocol to prevent dengue and yellow fever that include surveillance of neglected sylvatic vectors.

## Introduction

The extent of occurrence of both dengue and yellow fever, the most important arboviral diseases affecting humans worldwide^1,2^, has been changing due to intensive agriculture, irrigation, deforestation, population movements, rapid unplanned urbanisation, and phenomenal increases in international travel and trade^3^. The “urban” mosquito species *Aedes aegypti* and *Ae. albopictu*s are considered to be main vectors of these diseases, and so the shifting trend of both species’ ranges is of public health concern^4^. However, these mosquitoes are not the only species transmitting dengue and yellow fever, as some sylvatic species are also involved in viral spillover between primates^5^.

Haines^6^ and others in the 1980s suggested that global warming could become a driving factor for the future spread of mosquito-borne diseases. The incidence and distribution of these diseases are often conditioned by the abundance and distribution of their vectors ^7,8^. Numerous modelling studies have predicted range changes for *Ae. aegypti* and *Ae. albopictus* in response to climate change^9–11^. *Aedes* mosquito vectors could be currently expanding their distributions to more temperate climates across all continents where they now occur^12–14^, also favoured by globalisation through increased international travel and shipping. Many ports and airports are easy entry points for mosquito species such as *Ae. aegypti* and *Ae. albopictus*. For example, *Ae. albopictus* was introduced in Europe by the transport of tyres on ferries^15^. Predicting which might be the most favourable points of entry in the future could be useful for surveillance efforts. Recent research has suggested that, in the last 50 years of the 20^th^ century, the environmental conditions became around 1.5% more suitable for *Ae. aegypti* per decade globally, which could increase to 3.2–4.4% per decade by 2050^9^. Climate changes could even be forcing vectors to develop adaptation mechanisms facilitating infection spread^16^. However, up to now, the effects of climate change on sylvatic vectors have not been taken into account. Areas prone to sylvatic arboviral transmission to humans from non-human primates may be larger than estimated^13,17^. Anthropogenic disturbance in forests often allows the interaction of different mosquito communities with varying habitat preferences at ecotonal forest edges^18^. So, any attempt to make forecasting of dengue and yellow fever transmission risk in geographic contexts involving tropical areas should take both urban and sylvatic vectors into account. These species include *Ae. africanus*, *Ae. luteocephalus*, *Ae. vittatus* for dengue and yellow fever; *Sabethes chloropterus*, *Haemagogus leucocelaenus* and *Hg. janthinomys* for yellow fever and *Ae. niveus* for dengue^19^.

In this context, the objectives of this research are to answer the following questions: (1) Which mosquito species could be implicated in future increases in the risk of transmission for dengue and yellow fever due to climate change?, and (2) Where would these increases most likely occur?

## Methods

### Vector species

In order to analyse the effect of climate change on dengue and yellow fever vector distributions, we took into account the favourability models for these species published by Aliaga-Samanez et al.^13,17^ for the period 2001-2017, henceforth referred to as the baseline models. These models represent the degree of environmental favourability for vector species to occur on a grid of 18,874 hexagons of 7,774-km^2^ covering the whole World. Favourability is defined as the degree to which local conditions allow a higher or lower local probability than expected by chance (see below, equation 1). This chance probability is defined as the total prevalence of the analysed event^20^. Here, for each vector species, the total prevalence is the number of hexagons with presences divided by the number of hexagons in the study area. The baseline models considered urban vector species (*Ae. aegypti* and *Ae. albopictus)* and known sylvatic vector species (*Ae. africanus*, *Ae. luteocephalus*, *Ae. niveus*, *Ae. vittatus*, *Sabethes chloropterus*, *Haemagogus leucocelaenus* and *Hg. janthinomys*) (see Table S1).

### Forecasting the future distribution of vectors

In order to make forecasts for each of these species mentioned above, we recalculated favourability values (*F*) using the same equation as defined in the baseline models^13,17^, replacing the values of climate variables in each hexagon according to predictions by the Intergovernmental Panel on Climate Change (IPCC)^21^ (see below for details). Those baseline models^13,17^ are defined by the following equation:

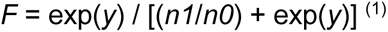

where *n1* and *n0* are, respectively, the number of presences and absences considered as dependent variables for model training, and *y* is a linear combination of predictor variables. The *y* equations for every species can be seen in Tables S2 and S3. Values for the non-climatic variables (i.e., Human Concentration, Infrastructures, Livestock, Topography, Agriculture, Ecosystem Types) forming part of the model were assumed not to change in the future period considered, thereby isolating the effect of a changing climate (Table S4). To consider uncertainties in our forecasts according to a range of variation in climatic predictions, we used climatic variable values from different climate change scenarios (CO_2_ representative concentration pathways, RCPs) and atmosphere–ocean general circulation models (GCMs) over two time periods, namely to 2041–2060 (“near future”) and to 2061–2080 (“distant future”). Five GCMs were selected. They provided predictions available for the whole World and with the lowest detected biases in the available GCMs with respect to actual climate data (Table S4)^22,23^: CESM1-CAM5, CNRM-CM5, FIO-ESM, GFDL-CM3, MPI-ESM-LR. We also considered two emissions scenarios: a stabilised emissions scenario RCP 4.5, and a high emissions scenario RCP 8.5. GCMs and RCP scenarios were taken from Climatologies at High resolution for the Earth Land Surface Areas (CHELSA) free high resolution climate data^24–27^. The basal model for each vector species was therefore projected to a total of 10 GCM-RCP scenario combinations per future period. Finally, a climate change forecast consensus was calculated using average favourability values of the 10 projections for a given period. For every disease and time period, a vector model projected to the future was got by combining the corresponding single-vector consensual forecasts with each other using the fuzzy logic operator “fuzzy union”^28^, which is equivalent to assigning the highest favourability value in each hexagonal unit.

### Expected rates of change in favourability

We mapped the expected increment of favourability by calculating the difference between forecasted favourability values (i.e., for 2041-2060 or 2061-2080) and those in the baseline model (2001-2017). In order to quantify to what extent the current favourability (*F_0_*) is modified globally in the future forecasts (*F_f_*), we calculated the fuzzy parameters of increment and maintenance according to the equations^21,29^:

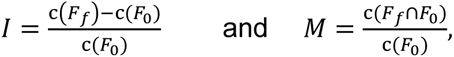

where *I* represents the global rate of increment in favourability, and *M* represents the global rate of maintenance of the original values. The factor c(*F_x_*) is the cardinality of the *F_x_* model or model projection – where favourability is treated as a fuzzy set^28^ – that is, the sum of all the hexagons’ favourability values (which, in turn, are treated as degrees of membership in the fuzzy set of hexagons favourable for the presence of vectors). The intersection between future (*F_f_*) and present (*F_0_*) favourability values, which is needed to calculate *M*, is defined as followed:

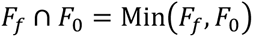

Positive increment values (*I*) indicate a net increase in favourability, that is, a gain of favourable areas; whereas negative values mean a net loss of favourable areas. The maintenance values (*M*) indicate the degree of overlap between present and forecasted favourable areas.

### Uncertainty in the forecasts

The local degree of uncertainty of forecasts resulting from a consensus based on averages (see above), was mapped by calculating the standard deviation, for each spatial unit, of the 10 possible favourability values projected for a given period (5 GCMs x 2 RCP scenarios). This standard deviation was interpreted as a measure of the reliability of forecasts in each hexagonal unit.

## Results

### *Ae. aegypti* and *Ae. albopictus* models projected into the future

Favourable areas for *Ae. aegypti*, according to the models projected into the future for the periods 2041–2060 and 2061–2080 show little differences compared to the 2001– 2017 baseline, with changes being perceptible only at regional scales (Figs. 1 and S1). The areas currently generally favourable for the presence of *Ae. aegypti* are expected to remain that way from now to 2080 (maintenance index value M, ≥ 0.96). In the near future (2041–2060), there could be a slight global net loss of favourability (increment index value, I = -0.03); whereas a positive increase (I = 0.075) is expected in the distant future (2061–2080). This increase would affect some regions of the Amazon basin and of Central Africa where favourability values, currently being intermediate-low, would turn into intermediate-high (Figs. 1 and S1). Extensive areas of the United States, Europe, and Australia could experience a favourability increase of 0.01 to 0.03 (Fig. S1) although these regions would remain generally unfavourable for the presence of *Ae. aegypti* (Fig. 1).

**Fig 1.**
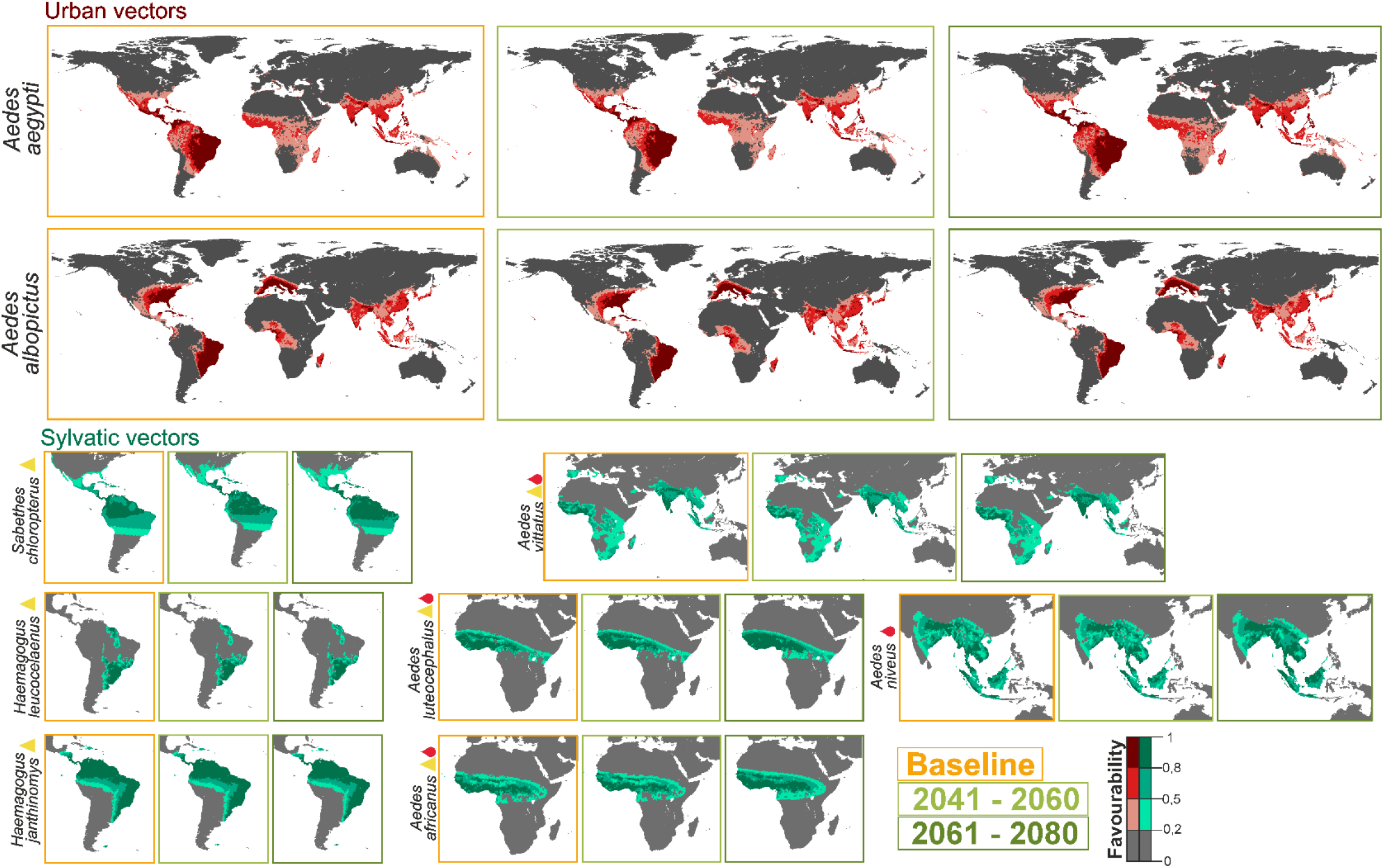
Urban and sylvatic vector model future projections for the periods 2041–2061 and 2061–2080. Models of the urban mosquitoes (*Ae. aegypti* and *Ae. albopictus*) and sylvatic mosquitoes (*H. janthinomys*, *H. leucocelaenus*, *S. chloropterus*, *Ae. luteocephalus*, *Ae. africanus*, *Ae. vittatus*, *Ae. niveus)* for the current time (2001–2017) and average model projections into the future for the periods 2041–2060 and 2061–2080. Yellow triangles represent yellow fever vectors and red drops represent dengue vectors.

Expected changes for *Ae. albopictus* are much lower (Fig. 1). Favourable areas for this species are expected to remain unchanged (M>0.99 in both future periods), while the global net gain in favourability could be slight, and would take place in the near future (I=0.019 for the period 2041–2060; I=0.020 for the period 2061–2080) (Fig. S1).

### Sylvatic vector models projected into the future

Changes predicted for sylvatic vectors depend on the mosquito species analyzed (Fig. 1). In America, favourable areas for *Haemagogus janthinomys* and *H. leucocelaenus* show no change with respect to the baseline. In contrast, high-favourability areas for *Sabethes chloropterus* in Central and South America seem to increase progressively north and southward. In Africa and Asia, changes could be negligible in the near future, whereas a little increase is expected in the distant future. This would be a north and southward increase in Central Africa for *Ae. luteocephalus* and for *Ae. africanus*; and a south and eastward increase in Asia for *Ae. niveus*. *Ae. vittatus*, which has populations in Europe, Africa and Asia, could experience a slight favourability increase elsewhere in the more distant future (Fig. 1).

### Combined (urban and sylvatic) vector models projected into the future

The models combining predictions for all the vectors of dengue show that areas currently favourable to the presence of these species will remain so in the near (2041–2060) and the distant (2061–2080) futures (both M>0.98). Despite favourability decreasing globally slightly in the near future (I=-0.006), favourability could increase locally by more than 7% in the north of West Africa and at the southern limits of the Himalayas (Figs. 2C and S2). However, this increase would not produce a noteworthy enlargement of high-favourability areas (compare Figs. 2A and 2B). In the more distant future, a 5% favourablity increase is expected (I=0.05), and so new areas could become favourable for the presence of dengue vectors in large areas of West and Central Africa and in South-East Asia, reaching Borneo (Figs. 2C and S2).

**Fig 2.**
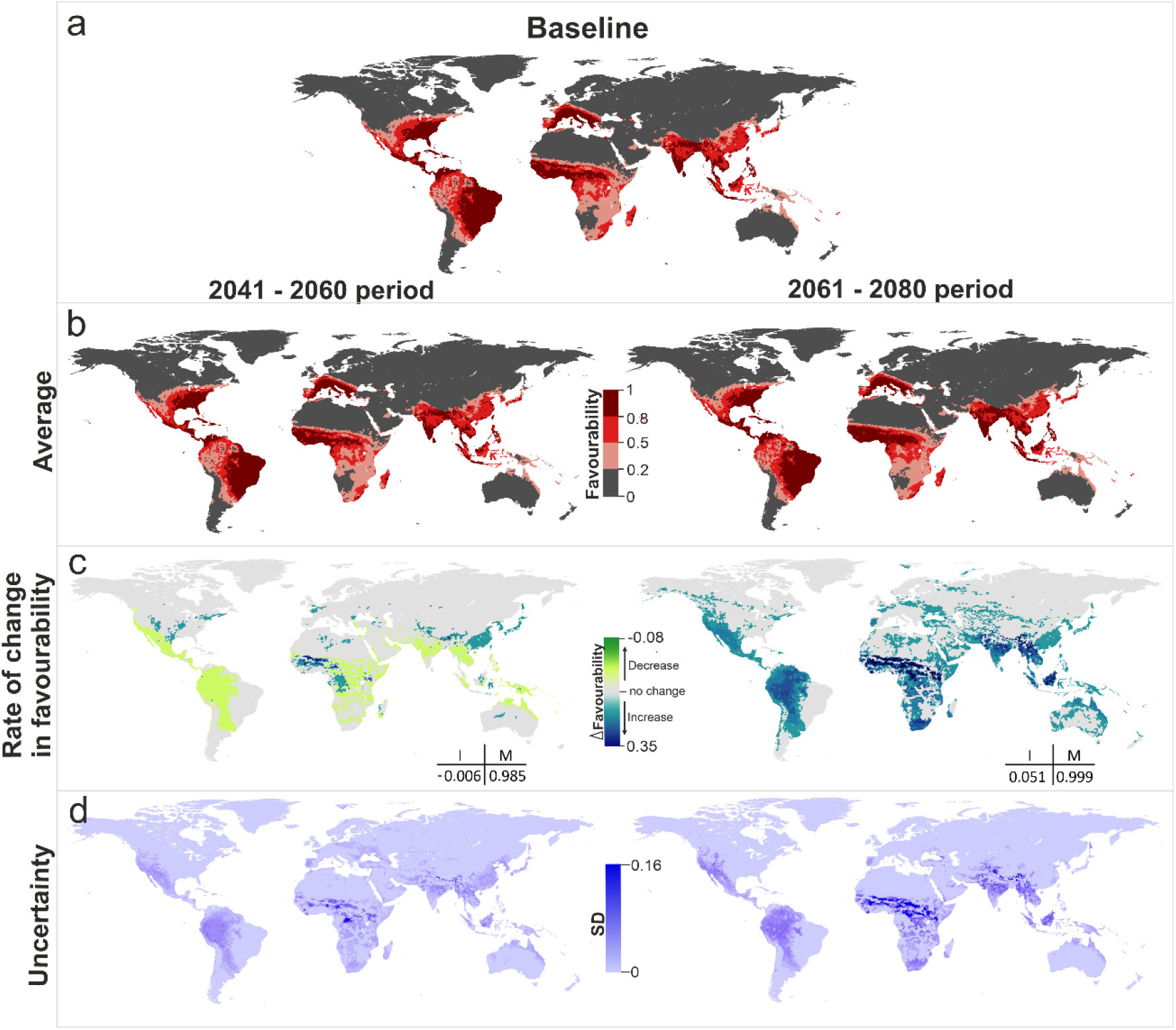
Dengue vector model projections into the future for the periods 2041–2060 and 2061– 2080. **a** Vector model for the current time (2001–2017), **b** average model projections into the future for the periods 2041–2060 and 2061–2080, **c** areas where favourability increases and decreases in the future relative to the present. Difference between the future projection and the current model. I: increment rate; M: maintenance rate. Positive values of I indicate a net increase in favourability, that is, a gain in favourable areas, whereas negative values of I mean a net loss of favourable areas. M indicates the degree to which the favourable areas in the current model overlap with the favourable forecasted areas. **d** Uncertainty of the vector model in the period 2041–2060 and 2061–2080. SD: Standard Deviation.

Combined predictions for yellow fever vectors (Fig. 3B) show a similar pattern to that of dengue. Favourable areas are predicted to remain unchanged in both future periods (M>0.97). Global favourability values could decrease slightly in the near future (I=-0.01), but might increase in the distant future (I=0.03). This increase is expected to affect principally the Amazon basin and West and Central Africa (Fig. 3C). The models indicate, instead, a regional favourability decrease of more than 0.01 points in coastal areas of Chile and Peru.

**Fig 3.**
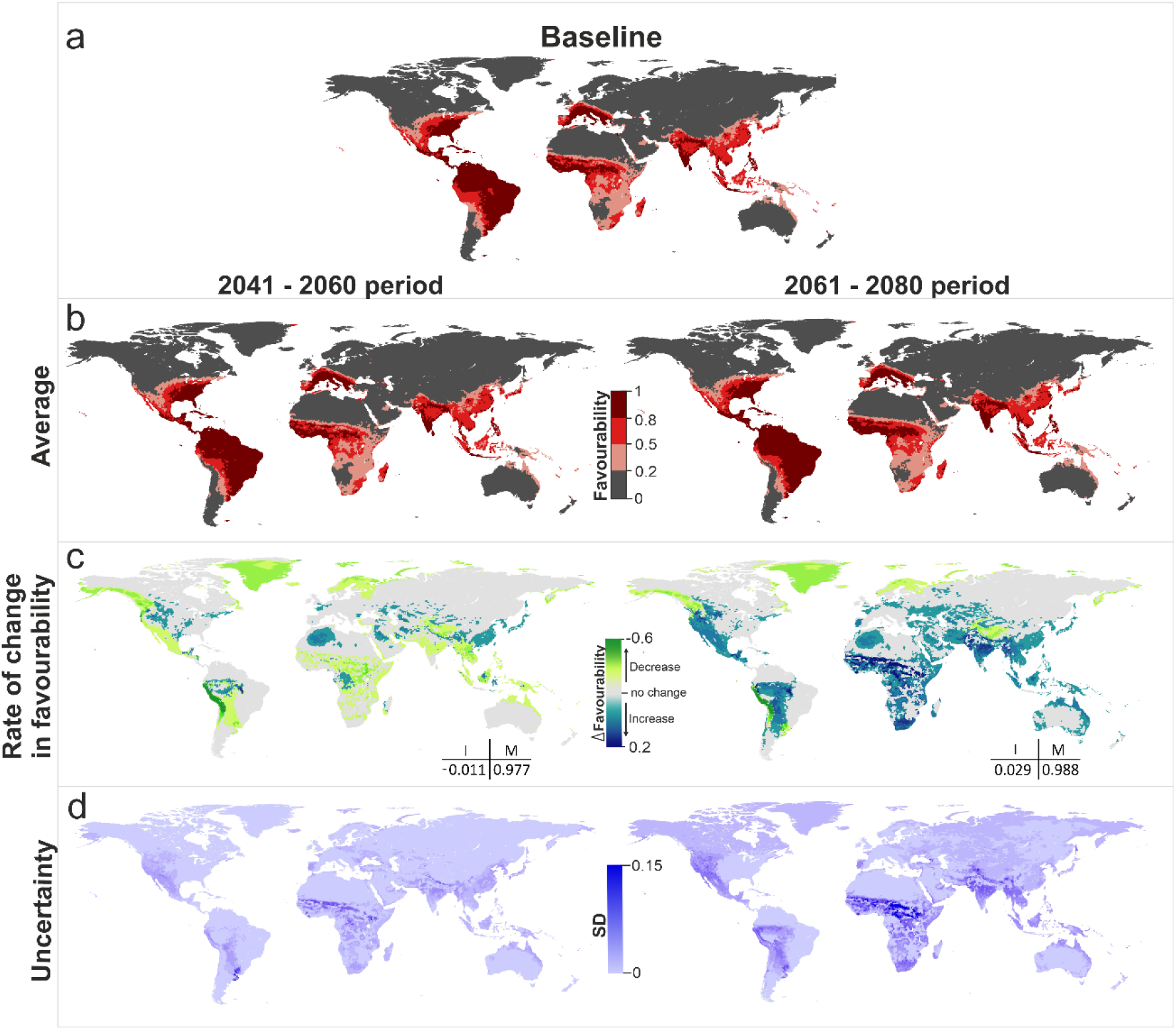
Yellow fever vector model future projections for the periods 2041–2060 and 2061– 2080. **a** Vector model for the current time (2001–2017), **b** average model projections into the future for the periods 2041–2060 and 2061–2080, **c** areas where favourability increases and decreases in the future relative to the present. Difference between the future projection and the current model. I: increment rate; M: maintenance rate. Positive values of I indicate a net increase in favourability, that is, a gain in favourable areas, whereas negative values of I mean a net loss of favourable areas. M indicates the degree to which the favourable areas in the current model overlap with the favourable forecasted areas. **d** Uncertainty of the vector model in the period 2041-2060 and 2061 – 2080. SD: Standard Deviation.

According to the uncertainty analysis, variations between scenario-model combinations in the near future mostly occur in the areas where average favourability is predicted to increase. In the more distant future, there is more consistency between projections, and high variations only occur in some areas of West and Central Africa (Figs. 2D and 3D). Some uncertainty is also observed at the southern limits of the Himalayas in predicted values regarding vectors transmitting the yellow fever virus.

## Discussion

This study is the first to analyze the effect of climate change on the distribution of all mosquito species transmitting dengue and yellow fever. Our projections for *Ae. albopictus* and *Ae. aegypti* predict that climate change could allow these species to expand into previously unoccupied temperate regions (Fig. 1). In addition, our projections have also detected species of sylvatic mosquitoes that may experience distribution changes, and could thus contribute to increasing disease transmission risk. This has implications for surveillance in the future.

The models show substantial heterogeneity in favourability patterns for the two primary vectors at finer scales. For example, *Ae. aegypti* has stable populations in all South American countries, while *Ae. albopictus* has not yet been detected in Peru, Ecuador, Chile, Bolivia and Uruguay, yet is already present in neighbouring countries. Aliaga-Samanez et al.^13^ describe an increase in favourable territories for both *Aedes* species, from the Brazilian coast to the Amazon basin during the last 20 years. According to our model projections, the Amazon basin could remain favourable for both species in the two future periods analysed. In particular, the Amazon could become even more favourable for *Ae. aegypti*, reaching intermediate-high values in Colombia, Peru and Brazil (Fig. 1). These results suggest that temperature changes might have already favoured the recent spread of *Aedes* species in the area, despite other studies predicting no new relevant variations in distribution under climate change in South America^30^. In North America, in contrast to Kraemer et al. (2019)^31^, who predict an expansion of *Ae. aegypti* between 2020 and 2050 in the United States, our projections are more conservative. We only predict a relevant increase in favourability for this species in the distant future, after 2060 (Fig. S1). Instead, both our projections and Kraemer and collaborators predict an increase in favourability for *Ae. albopictus* starting in the near future (Fig. S1).

In Asia, our models forecast an increase of favourability for *Ae. aegypti* in territories to the east and south of China (Fig. 1), which is consistent with Liu et al.^32^’s results. De Guilhem de Lataillade et al.^33^ observed that *Ae. aegypti* populations in Singapore, Taiwan, Thailand and New Caledonia are competent vectors for yellow fever transmission in that region of Asia and the Pacific, and our models also predict a slight increase in those countries, especially in Thailand (Fig. 1). Bonnin et al.^34^ found that *Ae. aegypti* and *Ae. albopictus* densities will increase in Southeast Asia due to future temperature increases. They also predict *Ae. aegypti* densities will increase from 25% (with climate mitigation measures) to 46% without, with smaller increases of 13% and 21% for *Ae. albopictus*. On Reunion Island, our projections predict a slight increase in favourability for *Ae. albopictus* in both future time periods, agreeing with Lamy et al.^35^ who found that *Ae. albopictus* abundance will increase in 2070–2100. Central Africa could also see a spread of favourable areas for *Ae. aegypti* (Fig. 1). Gaythorpe et al. ^36^ report that Central Africa Republic is one of the countries most likely to see an increase in yellow fever transmission. Unfortunately, Africa lacks uniform surveillance policies ^37^. The results of this work could therefore be used to highlight territories where surveillance efforts should be focused. In Europe, in the distant future, our projections predict that *Ae. aegypti* could find favourable areas in Spain, the Netherlands and Portugal and, more intensively, in Italy, Turkey and Greece (Fig. S1). This agrees with Kraemer et al.’s ^31^ suggestion that, around 2080, this species could become established in Italy and Turkey. *Ae. albopictus* is already established in many European countries and will surely continue to find favourable areas in the future. Favourability is predicted to increase after 2040 in central Europe, in countries such as France, Germany, Belgium and United Kingdom. Kraemer et al.^31^ also forecasted *Ae. albopictus* to spread widely across Europe, reaching large parts of France and Germany.

Focusing on the response of sylvatic mosquitoes to climate changes (Figs. 1, 2 and 3), *Ae. niveus* and *Ae. vittatus* should be subject to survey as dengue vectors in Asia, where areas favourable to their presence could spread in the hinterlands of India, in the south-east of China, and in the South-Asian countries, especially in Borneo. In west and central sub-Saharan Africa, according to our results, attention should be paid to the three sylvatic vectors of dengue and yellow fever, *Ae. vittatus*, *Ae. luteocephalus* and *Ae. africanus*, especially in Cameroon, Central Africa Republic and north of Democratic Republic of Congo. However, the current entomological capacity in Africa is primarily focused on malaria vectors, and most countries lack routine surveillance programmes, trained personnel and control activities focusing on *Aedes* and the viruses they transmit^38^. As outbreaks of *Aedes*-borne arboviruses continue to increase in Africa, it would be critical to establish a solid public health entomology infrastructure for *Aedes* mosquitoes to contain and prevent further outbreaks^39^.

Both in Africa and in Asia, the most relevant changes are forecasted for the distant future (2061-2080), but in South America, the increase in favourability for the yellow fever vector *Sabetes chloropterus*, forecasted for the jungle areas of Bolivia, Peru and Brazil, could start in the near future. The climatic conditions could become, however, less favourable for this species in western Peru and Bolivia. On the other hand, our models suggest no change in favourable zones for *H*. *janthinomys*. Sadeghieh et al.^40^ suggest that many areas in Brazil will become unsuitable for *Haemagogus* spp. to survive and suboptimal for yellow fever virus replication under climate change. We therefore suggest that surveillance and prevention measures planned by the Pan American Health Organization (PAHO) for urban mosquitoes also include specifications and survey planning for *S. chloropterus*. In Brazil, sylvatic yellow fever outbreaks that have occurred since 2017 highlight the need to strengthen surveillance for zoonotic yellow fever in non-human primates^17,41^.

Such variation has implications for how climate change may influence establishment success in areas currently absent from urban mosquito species. For example, taking into account that some of the most important ports in Europe are located in Barcelona, Rotterdam, Valencia, and Antwerp^42^, these could be important points for establishment of *Ae. aegypti* in case of accidental entry. There is already evidence of accidental arrivals of this mosquito species in northern Europe. For instance, in 2010 national surveillance in the Netherlands detected the presence of *Ae. aegypti* in a shipment of tires from Miami ^43,44^, and in 2016, *Ae. aegypti* was again detected in the Netherlands at the Schiphol International airport^45^. Da Re et al.^46^ assessed the probability that *Ae. aegypti*, which currently occurs just on the east coast of the Black Sea, Turkey, Bulgaria and Russia^47^, spread through continental Europe. They selected five European ports taking into account environmental conditions and the economic importance of the harbours, and suggested that the city most likely to be at risk of establishment was Barcelona, followed by Algeciras. Our results concur. In Europe our model suggests that Barcelona, Algeciras, Venice, Sardinia, Antwerp, Naples and the west coast of Portugal are favourable areas (0.2<F<0,5) for the presence of *Ae. aegypti* (Fig. 1). In addition, we detect areas with intermediate-high favourability values (0.5<F<0.8) in Valencia, Rotterdam, Istanbul and Naples (Fig. 1). For *Ae. albopictus,* favourable conditions are predicted to occur in the 7 countries where it is not yet established, which will continue to be favourable through to 2080 (Fig. 1).

For *Ae. aegypti,* re-introduction in Europe is possible with many pathways and entry points. The question is whether, once the introduction has happened, environmental conditions are suitable for the species to establish successfully. Although Da Re et al.^46^ detected that the areas of Genoa, Venice and Rotterdam are not suitable for establishment of *Ae. aegypti* populations now, they suggested that climate change could create conditions for a future invasion. Both Campbell et al.^7^ and Ryan et al.^10^ concur that the potential distribution of *Ae. aegypti* under future conditions could increase in most of Europe, even reaching the north of the continent as a consequence of climate change. However, our model projections for 2060 and 2080 continue to predict the same favourable areas that are currently suitable. According to our models, the risk of *Ae. aegypti* new introduction in Europe would remain concentrated in cities that are equipped with large harbours and/or airports, most probably favoured by the chances of arrival than by climate changes.

Similarly, the introduction of *Ae. aegypti* in the Canary Islands, northwest of Africa, has so far not lead to establishment. In early 2022, *Ae. aegypti* larvae were detected in La Palma, one of the Canary Islands, but extensive entomological surveillance work has not detected any life-stage since^48^. Also in 2022, this species was detected on another of the Canary Islands, in Tenerife, but reinforced entomological surveillance has not detected further occurrence there^49^. The high level of regular communication with nearby endemic regions such as Madeira Island (Portugal) and, to a lesser extent, with the archipelago of the Republic of Cape Verde, means that there is a real risk of introduction of the vector to the islands^49^.

In South America, *Ae. aegypti* has been introduced and established in new regions of Peru in recent years, and also at new altitudes. This happened in Piura (North of the country) and in Huanuco (Centre-East), at 1,959 and 2,227 metres above sea level, respectively^50^. In 2022, the number of dengue cases in Peru exceeded that of the previous year^51^. Our models predict an increase in favourability for *Ae. aegypti* in different regions of Peru for both future periods (Fig. 1), which could increase the number of dengue cases in the following years. In this scenario, the Pan American Health Organization (PAHO) has established, since 2019, a manual for indoor residual spraying in urban areas for the control of *Ae. aegypti* in American regions^41^.

The situation for *Ae. albopictus* is different. The establishment of *Ae. albopictus* in western South America, in countries such as Peru, could further favour the risk of dengue and yellow fever transmission, in addition to favouring the link between the urban and jungle cycle, since *Ae. albopictus* acts as a "bridge vector". Our models do not predict a change in the favourable zones with respect to the baseline model, predicting that it will continue to be favourable in both future periods in southeastern Peru (Fig. 1). Peru has reinforced surveillance measures on the borders with Brazil and Colombia, has developed a technical standard for entomological surveillance and control of *Ae. aegypti* and surveillance of the entry of *Ae. albopictus* into the national territory^52^. In Ecuador, this species was reported for the first time in 2017 in Guayaquil, and is currently expanding its distribution in the country^53,54^. Guayaquil is a coastal city and has the main port in the country, responsible for more than 80% of exports, and vulnerable to invasive species^54^. In Australia, our models predict that *Ae. albopictus* will find favourable areas in eastern Australia in the future (Fig. 1), agreeing with Laporta et al. ^55^. However, the Australian government has had a well-structured control plan in place since 2005 which has resulted in the successful suppression of *Ae. albopictus* populations at major transport hubs in the Torres Strait^56^. In Europe, this species has established stable populations since 1979, probably introduced via the Durres Port (Albania) through the entry of goods from China^57^. *Ae. albopictus* has now been introduced in 33 European countries and is established in 26. This species might have caused dengue outbreaks already in France, Italy and Croatia and autochthonous cases in Spain^58,59^. In 2022, an increase in imported cases of dengue from Cuba was detected in Europe due to limited fumigation in Cuba ^60^, and, at the beginning of 2023, autochthonous cases were detected for the first time in Ibiza (Balearic Islands).

According to the WHO, funding is needed in south-eastern Asia for strengthening surveillance, reporting and vector management ^61^, as a response to the increased burden of dengue experienced since 2011^62^, and although no autochthonous yellow fever cases have been reported in the Asia-Pacific regions, surveillance for imported cases is recommended. The combination of repeated introductions of imported cases, an immunologically naïve local population in an environment suitable for transmission, and climate change that will continue to favour its presence in the future could increase the risk of yellow fever occurrence in Asia.

## Conclusions

Climate change could modify and increase the risk of zoonotic diseases through its effects on vector configuration and distribution. Increased favourability for urban and sylvatic mosquitoes could contribute to the expansion of risk areas for dengue and yellow fever transmission. Our study suggests such increases could occur particularly in Thailand, Peru, Colombia, Brazil, China, India, Central African Republic, Spain, Italy, Netherlands, and Portugal. *Ae. aegypti*, *Ae. albopictus*, *Ae. vittatus*, *Ae. luteochephalus*, *Ae. africanus*, *Ae. niveus* and *S. chloropterus,* according to our analysis, will find new favourable areas or favourability will increase in currently occupied areas. Our results suggest the need to establish a surveillance protocol to focus and prevent new cases of dengue and yellow fever that includes neglected sylvatic vectors.

## Availability of data and materials

All the data relevant generated or analysed during this study are included in this published article and its Supplemental Material.

## Author contributions

AA-S, RD, RR and OJ designed and implemented the study. AA-S and OJ managed the curation of the database and analysed the data. AA-S and performed and applied the modelling procedures. AA-S, RD and OJ interpreted the data, wrote, reviewed, and edited the manuscript. SM interpreted and reviewed about the health interpretation of the main results. MK, RR and OJ revised the different versions of the manuscript. RR and OJ managed the funding acquisition and project administration. All authors read and approved the final version of the manuscript.

## References

1. Kuno, G. A Re-Examination of the History of Etiologic Confusion between Dengue and Chikungunya. PLoS Negl. Trop. Dis. 9, 1–11 (2015).

2. Colón-González, F. J. et al. Projecting the risk of mosquito-borne diseases in a warmer and more populated world: a multi-model, multi-scenario intercomparison modelling study. Lancet Planet. Heal. 5, e404–e414 (2021).

3. WHO. A global brief on vector-borne diseases. https://apps.who.int/iris/handle/10665/111008 (World Health Organization Press, 2014).

4. Rupasinghe, R., Chomel, B. B. & Martínez-López, B. Climate change and zoonoses: A review of the current status, knowledge gaps, and future trends. Acta Trop. 226, 106225 (2022).

5. Hanley, K. A. et al. Fever versus fever: The role of host and vector susceptibility and interspecific competition in shaping the current and future distributions of the sylvatic cycles of dengue virus and yellow fever virus. Infect. Genet. Evol. 19, 292–311 (2013).

6. Haines, A. Global warming and health. BMJ 302, 669–670 (1991).

7. Campbell-Lendrum, D., Manga, L., Bagayoko, M. & Sommerfeld, J. Climate change and vector-borne diseases: what are the implications for public health research and policy? Philos. Trans. R. Soc. B Biol. Sci. 370, 1–8 (2015).

8. Messina, J. P. et al. The many projected futures of dengue. Nat. Rev. Microbiol. 2015 134 13, 230–239 (2015).

9. Iwamura, T., Guzman-Holst, A. & Murray, K. A. Accelerating invasion potential of disease vector Aedes aegypti under climate change. Nat. Commun. 2020 111 11, 1–10 (2020).

10. Ryan, S. J., Carlson, C. J., Mordecai, E. A. & Johnson, L. R. Global expansion and redistribution of Aedes-borne virus transmission risk with climate change. PLoS Negl. Trop. Dis. 13, e0007213 (2018).

11. Campbell, L. P. et al. Climate change influences on global distributions of dengue and chikungunya virus vectors. Philos. Trans. R. Soc. B Biol. Sci. 370, 1–9 (2015).

12. Watts, N. et al. The 2020 report of The Lancet Countdown on health and climate change: responding to converging crises. Lancet 397, 129–170 (2021).

13. Aliaga-Samanez, A. et al. Worldwide dynamic biogeography of zoonotic and anthroponotic dengue. PLoS Negl. Trop. Dis. 15, e0009496 (2021).

14. Khan, S. U. et al. Current and Projected Distributions of Aedes aegypti and Ae. albopictus in Canada and the U.S. Environ. Health Perspect. 128, (2020).

15. Marrama Rakotoarivony, L. & Schaffner, F. ECDC guidelines for the surveillance of invasive mosquitoes in Europe. Eurosurveillance vol. 17 (2012).

16. Filho, W. L., Ternova, L., Parasnis, S. A., Kovaleva, M. & Nagy, G. J. Climate Change and Zoonoses: A Review of Concepts, Definitions, and Bibliometrics. Int. J. Environ. Res. Public Heal. 2022 19, 893 (2022).

17. Aliaga-Samanez, A., Real, R., Segura, M., Marfil-Daza, C. & Olivero, J. Yellow fever surveillance suggests zoonotic and anthroponotic emergent potential. *Commun*. Biol. 5, (2022).

18. Meyer Steiger, D. B., Ritchie, S. A. & Laurance, S. G.W. Mosquito communities and disease risk influenced by land use change and seasonality in the Australian tropics. Parasites and Vectors 9, 1–13 (2016).

19. Carvalho, B. M., Rangel, E. F. & Vale, M. M. Evaluation of the impacts of climate change on disease vectors through ecological niche modelling. Bull. Entomol. Res. 107, 419–430 (2017).

20. Acevedo, P. & Real, R. Favourability: Concept, distinctive characteristics and potential usefulness. Naturwissenschaften 99, 515–522 (2012).

21. Romero, D., Olivero, J., Márquez, A. L., Báez, J. C. & Real, R. Uncertainty in distribution forecasts caused by taxonomic ambiguity under climate change scenarios: a case study with two newt species in mainland Spain. J. Biogeogr. 41, 111–121 (2014).

22. McSweeney, C. F., Jones, R. G., Lee, R. W. & Rowell, D. P. Selecting CMIP5 GCMs for downscaling over multiple regions. Clim. Dyn. 44, 3237–3260 (2015).

23. Sanderson, B. M., Knutti, R. & Caldwell, P. A representative democracy to reduce interdependency in a multimodel ensemble. J. Clim. 28, 5171–5194 (2015).

24. Nikolaus Karger, D., Schmatz, D. R., Dettling, G. & Zimmermann, N. E. High-resolution monthly precipitation and temperature time series from 2006 to 2100. Sci. Data 7, 248 (2020).

25. Karger, D. N., Wilson, A. M., Mahony, C., Zimmermann, N. E. & Jetz, W. Global daily 1 km land surface precipitation based on cloud cover-informed downscaling. Sci. data 8, 307 (2021).

26. Karger, D. N. et al. Climatologies at high resolution for the earth’s land surface areas. Sci. Data 4, 1–20 (2017).

27. Knutti, R., Masson, D. & Gettelman, A. Climate model genealogy: Generation CMIP5 and how we got there. Geophys. Res. Lett. 40, 1194–1199 (2013).

28. Estrada, A., Real, R. & Vargas, J. M. Using crisp and fuzzy modelling to identify favourability hotspots useful to perform gap analysis. Biodivers. Conserv. 17, 857–871 (2008).

29. Kuncheva, L. I. Using measures of similarity and inclusion for multiple classiÿer fusion by decision templates. Fuzzy Sets Syst. 122, 401–407 (2001).

30. Kamal, M., Kenawy, M. A., Rady, M. H., Khaled, A. S. & Samy, A. M. Mapping the global potential distributions of two arboviral vectors Aedes aegypti and Ae. albopictus under changing climate. PLoS One 13, e0210122 (2018).

31. Kraemer, M. U. G. et al. Past and future spread of the arbovirus vectors *Aedes aegypti* and *Aedes albopictus*. Nat. Microbiol. 4, 854–863 (2019).

32. Liu, B. et al. Modeling the present and future distribution of arbovirus vectors Aedes aegypti and Aedes albopictus under climate change scenarios in Mainland China. Sci. Total Environ. 664, 203–214 (2019).

33. de Guilhem de Lataillade, L., et al. Risk of yellow fever virus transmission in the Asia-Pacific region. Nat. Commun. 2020 111 11, 1–10 (2020).

34. Bonnin, L. et al. Predicting the Effects of Climate Change on Dengue Vector Densities in Southeast Asia through Process-Based Modeling. Environ. Health Perspect. 130, (2022).

35. Lamy, K., Tran, A., Portafaix, T., Leroux, M. D. & Baldet, T. Impact of regional climate change on the mosquito vector Aedes albopictus in a tropical island environment: La Réunion. Sci. Total Environ. 875, 162484 (2023).

36. Gaythorpe, K. A. M., Hamlet, A., Cibrelus, L., Garske, T. & Ferguson, N. M. The effect of climate change on yellow fever disease burden in Africa. Elife 9, 1–27 (2020).

37. Weetman, D. et al. Aedes Mosquitoes and Aedes-Borne Arboviruses in Africa: Current and Future Threats. Int. J. Environ. Res. Public Health 15, 220 (2018).

38. Buchwald, A. G., Hayden, M. H., Dadzie, S. K., Paull, S. H. & Carlton, E. J. Aedes-borne disease outbreaks in West Africa: A call for enhanced surveillance. Acta Trop. 209, 105468 (2020).

39. Dadzie, S. K. et al. Building the capacity of West African countries in Aedes surveillance: inaugural meeting of the West African Aedes Surveillance Network (WAASuN). Parasit. Vectors 15, 381 (2022).

40. Sadeghieh, T. et al. Yellow fever virus outbreak in Brazil under current and future climate. Infect. Dis. Model. 6, 664–677 (2021).

41. PAHO. Plan de acción sobre entomología y control de vectores 2018-2023 (56 Consejo Directivo). https://www.paho.org/es/documentos/cd5611-plan-accion-sobre-entomologia-control-vectores-2018-2023 (2019).

42. Eurostat. Top 20 EU ports handling containers, 2010, 2019 and 2020 (million TEUs). https://ec.europa.eu/eurostat/statistics-explained/index.php?title=File:Top_20_EU_ports_handling_containers,_2010,_2019_and_2020_(million_TEUs)_V2.png Accessed 2022-06-28 https://ec.europa.eu/eurostat/statistics-explained/index.php?title=File:Top_20_EU_ports_handling_containers,_2010,_2019_and_2020_(million_TEUs)_V2.png (2021).

43. Scholte, E. J. et al. Introduction and control of three invasive mosquito species in the Netherlands, July-OctOber 2010. Eurosurveillance 15, 1–4 (2010).

44. Brown, J. E. et al. Aedes aegypti Mosquitoes Imported into the Netherlands, 2010. Emerg. Infect. Dis. 17, 2335 (2011).

45. Ibañez-Justicia, A. et al. The first detected airline introductions of yellow fever mosquitoes (*Aedes aegypti*) to Europe, at Schiphol International airport, the Netherlands. Parasites and Vectors 10, 1–9 (2017).

46. Da Re, D., Montecino-Latorre, D., Vanwambeke, S. O. & Marcantonio, M. Will the yellow fever mosquito colonise Europe? Assessing the re-introduction of *Aedes aegypti* using a process-based population dynamical model. Ecol. Inform. 61, 101180 (2021).

47. ECDC. *Aedes aegypti* - current known distribution: March 2022. https://www.ecdc.europa.eu/en/publications-data/aedes-aegypti-current-known-distribution-march-2022 accessed 2022-05-07 https://www.ecdc.europa.eu/en/publications-data/aedes-aegypti-current-known-distribution-march-2022 (2022).

48. Centro de Coordinación de Alertas y Emergencias Sanitarias del Ministerio de Sanidad de España. Evaluación rápida de riesgo: Identificación del mosquito Aedes aegypti en la isla de La Palma. (2022).

49. Centro de Coordinación de Alertas Y Emergencias Sanitarias del Ministerio de Sanidad de España. Evaluación rápida de riesgo: Identificación del mosquito Aedes aegypti en Santa Cruz de Tenerife. https://www.mscbs.gob.es/profesionales/saludPublica/ccayes/alertasActual/docs/20171226_Aedes-aegypti_en_Fuerteventura_ERR.pdf (2023).

50. More, M., Castañeda, C. & Suyón, M. NEW ALTITUDINAL REGISTRATION OF Aedes aegypti IN THE REGION OF PIURA, PERU. Rev Peru Med Exp Salud Publica 35, 536–537 (2018).

51. Ministerio de Salud de Perú. Minsa: Más de 58 mil casos de dengue se han notificado en regiones del país en 2022. https://www.gob.pe/institucion/minsa/noticias/653797-minsa-mas-de-58-mil-casos-de-dengue-se-han-notificado-en-regiones-del-pais-en-2022 (2022).

52. DIGESA. Fronteras seguras para prevenir el ingreso del mosquito ‘Aedes albopictus’, transmisor de chikungunya, en el Perú. (2023).

53. Carrazco-Montalvo, A. et al. Establishment, Genetic Diversity, and Habitat Suitability of Aedes albopictus Populations from Ecuador. Insects 13, (2022).

54. Ponce, P., Morales, Di., Argoti, A. & Cevallos, V. E. First Report of Aedes (Stegomyia) albopictus (Skuse) (Diptera: Culicidae), the Asian Tiger Mosquito, in Ecuador. J. Med. Entomol. 55, 248 (2018).

55. Laporta, G. Z. et al. Global Distribution of Aedes aegypti and Aedes albopictus in a Climate Change Scenario of Regional Rivalry. Insects 14, (2023).

56. Muzari, M. O. et al. Holding back the tiger: Successful control program protects Australia from Aedes albopictus expansion. PLoS Negl. Trop. Dis. 11, e0005286 (2017).

57. Adhami, J. & Reiter, P. Introduction and establishment of Aedes (Stegomyia) albopictus skuse (Diptera: Culicidae) in Albania. J. Am. Mosq. Control Assoc. 14, 340–343 (1998).

58. Gossner, C. M., Ducheyne, E. & Schaffner, F. Increased risk for autochthonous vector-borne infections transmitted by *Aedes albopictus* in continental europe. Eurosurveillance 23, 2–7 (2018).

59. Centro de Coordinación de Alertas y Emergencias Sanitarias. Ministerio de Sanidad Consumo y Bienestar. Dengue autóctono en España 2a actualización. Evaluación rápida de riesgo. https://www.mscbs.gob.es/profesionales/saludPublica/ccayes/analisisituacion/doc/ERR_Dengue_autoctono_mayo2019.pdf (2019).

60. EL Pais. España: Los hospitales españoles detectan un fuerte incremento de casos de dengue en turistas procedentes de Cuba | Sociedad | EL PAÍS. (2022).

61. Nagpal, B. et al. Strengthening of vector control in South-East Asia: Outcomes from a WHO regional workshop. J. Vector Borne Dis. 55, 247 (2018).

62. Bangert, M., et al. Economic analysis of dengue prevention and case management in the Maldives. PLoS Negl. Trop. Dis. 12, e0006796 (2018).

