## Supplementary Information for "Potential climate change effects on the distribution of urban and sylvatic dengue and yellow fever vectors"

### Table of Contents

**- Table S1**. Sources of dengue and yellow fever vector records in Aliaga-Samanez et al. (2021, 2022) (species considered: *Aedes aegypti*, *Aedes albopictus*, *Haemagogus leucocelaenus*, *Haemagogus janthinomys*, *Sabethes chloropterus*, *Aedes luteocephalus*, *Aedes africanus*, *Aedes vittatus* and *Aedes niveus*).

**- Table S2.** Resulting logit functions of the baseline models in Aliaga-Samanez *et al*. 2021, 2022. Logit functions are linear combinations (*y*) of predictor variables that form part of the favourability (*F*) function: $F = \exp(y) / [(n1/n0) + \exp(y)]$. In the case of the *Ae. aegypti* and *Ae. albopictus* vectors, the predictor variable "*Y-20$^{th}$ century*" was included, which is the logit equation of the 20$^{th}$ century model (see Aliaga-Samanez et al. 2021, 2022). This predictor variable is detailed in table S3. B: variable coefficient; SE: standard error; W: Wald parameter; DF: degrees of freedom; S: statistical significance.

- **Table S3.** Logit equations of the vector models (*Aedes aegypti* and *Aedes albopictus*) of the 20$^{th}$ century (i.e., linear combinations of predictor variables that form part of the logistic-regression equations). B: variable coefficient; SE: standard error; W: Wald parameter; DF: degrees of freedom; S: statistical significance.

**-Table S4.** Predictor variables considered for baseline model training in Aliaga-Samanez et al. (2021, 2022).

**- Fig. S1**. Areas where favourability increases and decreases in the future relative to the present. Difference between the future projection and the current model. I: increment rate; B: maintenance rate (Romero et al. Journal of Biogeography 41.1 (2014): 111-121). Positive values of I indicate a net increase in favourability, that is, a gain in favourable areas, whereas negative values of I mean a net loss of favourable areas. M indicates the degree to which the favourable areas in the current model overlap with the favourable forecasted area.

**- Fig. S2**. Rate of change in favourability of dengue and yellow fever vectors models. Zoomed details of areas where favourability increases and decreases in the future relative (2041-2060 and 2061 – 2080) to the present.

**Table S1.** Sources of dengue and yellow fever vector records in Aliaga-Samanez *et al*. 2021, 2022 (species considered: *Aedes aegypti, Aedes albopictus, Haemagogus leucocelaenus, Haemagogus janthinomys, Sabethes chloropterus, Aedes luteocephalus, Aedes africanus, Aedes vittatus* and *Aedes niveus*).


1. Kraemer MUG, Sinka ME, Duda KA, Mylne AQN, Shearer FM, Barker CM, et al. The global distribution of the arbovirus vectors *Aedes aegypti* and *Ae. albopictus*. Elife. 2015;4: e08347.
2. Mosquito alert. Ciencia ciudadana para investigar y controlar mosquitos transmisores de enfermedades. 2021. http://www.mosquitoalert.com/. Accessed 11 July 2021
3. Vectorbase. Bioinformatics Resources for Invertebrate Vectors of Human Pathogens. 2021. https://vectorbase.org/vectorbase/app. Accessed 07 May 2021
4. de Abreu FVS, Ribeiro IP, Ferreira-de-Brito A, Santos AAC dos, de Miranda RM, Bonelly I de S, et al. *Haemagogus leucocelaenus* and *Haemagogus janthinomys* are the primary vectors in the major yellow fever outbreak in Brazil, 2016–2018. Emerg Microbes Infect. 2019; 8:218-231.
5. WHO. Geographical distribution of arthropod-borne diseases and their principal vectors [Internet]. Geneva, 1989, Switzerland: World Health Organization Press.
6. GBIF. Free and open access to biodiversity data. 2021. https://www.gbif.org/. Accessed 11 July 2021
7. VectorMap. Resource for collection records and other geospatial data relating to arthropods of biomedical importance. 2021. https://vectormap.si.edu/. Accessed 29 May 2021
8. Childs ML, Nova N, Colvin J, Mordecai EA. Mosquito and primate ecology predict human risk of yellow fever virus spillover in Brazil. Philos Trans R Soc B Biol Sci. Royal Society Publishing; 2019;374.
9. Ribeiro ALM, Miyazaki RD, Silva M, Zeilhofer P. Spatial and temporal abundance of three sylvatic yellow fever vectors in the influence area of the Manso hydroelectric power plant, Mato Grosso, Brazil. J Med Entomol. 2012;49:225–6.
10. Pajot F-X, Geoffroy B, Chippaux JP. Écologie Dyar d ' *Haemagogus janthinomys* données. 1985;XXIII:209–16.
11. Diagne CT, Diallo D, Faye O, Ba Y, Faye O, Gaye A, et al. Potential of selected Senegalese *Aedes spp*. mosquitoes (Diptera: Culicidae) to transmit Zika virus. BMC Infect Dis. 2015;15: 2–7. doi:10.1186/s12879-015-1231-2
12. Joseph AO, Adepeju S-OI, Omosalewa OB. Distribution, abundance and diversity of mosquitoes in Akure, Ondo State, Nigeria. J Parasitol Vector Biol. 2013;5: 132–136. doi:10.5897/JPVB2013.0133
13. Diallo D, Sall AA, Buenemann M, Chen R, Faye O, Diagne CT, et al. Landscape Ecology of Sylvatic Chikungunya Virus and Mosquito Vectors in Southeastern Senegal. PLoS Negl Trop Dis. 2012;6: 1–14. doi:10.1371/journal.pntd.0001649
14. Diallo M, Thonnon J, Traore-Lamizana M, Fontenille D. Vectors of Chikungunya virus in Senegal: Current data and transmission cycles. Am J Trop Med Hyg. 1999;60: 281–286. doi:10.4269/ajtmh.1999.60.281
15. Bang YH, Bown DN, Arata AA. Ecological studies on *Aedes Africanus* (Diptera: Culicidae) and associated species in Southeastern Nigeria. J Med Entomol. 1980;17: 411–416. doi:10.1093/jmedent/17.5.411
16. Hervy JP, Legros F, Roche JC, Monteny N, Diaco B. Circulation du virus Dengue 2 dans plusieurs milieux boisés des savanes soudaniennes de la région de Bobo-Dioulasso (Burkina Faso). Cah ORSTOM Ser Ent Med Parasitol. 1984;22: 135–143.
17. Roche JC, Cordellier R, Hervy JP. Ninety-six strains of dengue 2 virus isolated from mosquitoes collected in Ivory Coast and Upper Volta. Ann Virol. 1983;134: 233–244. Available: http://www.embase.com/search/results?subaction=viewrecord&from=export&id=L13088076%5Cnhttp://sfx.hul.harvard.edu/sfx_local?sid=EMBASE&issn=02425017&id=doi:&atitle=Ninety-six+strains+of+dengue+2+virus+isolated+from+mosquitoes+collected+in+Ivory+Coast+and+U
18. Onyido A, Ezike V, Ozumba N, Nwosu E, Ikpeze O, Obiukwu M, et al. Crepuscular Man-Biting Mosquitoes Of A Tropical Zoological Garden In Enugu, South-Eastern Nigeria. Internet J Parasit Dis. 2012;4: 4–9. doi:10.5580/11a2
19. Agwu EJ, Igbinosa IB, Isaac C. Entomological assessment of yellow fever-epidemic risk indices in Benue State, Nigeria, 2010-2011. Acta Trop. 2016;161: 18–25. doi:10.1016/j.actatropica.2016.05.005
20. Ahmed UA, Sani Z. Studies of mosquitoes in Hadejia Emirate, Jigawa state, Nigeria. Book of Proceedings of the Academic Conference on Positioning Sub-Sahara Africa for Development in the new Development. 2016. pp. 1–6.
21. Bang YH, Bown DN, Onwubiko AO. Prevalence of larvae of potential yellow fever vectors in domestic water containers in south-east Nigeria. Bull World Health Organ. 1981;59: 107–114.
22. Diallo M, Ba Y, Sall AA, Diop OM, Ndione JA, Mondo M, et al. Amplification of the sylvatic cycle of dengue virus type 2, Senegal, 1999-2000: Entomologic findings and epidemiologic considerations. Emerg Infect Dis. 2003;9: 362–367. doi:10.3201/eid0903.020219
23. Anosike JC, Nwoke BEB, Okere AN, Oku EE, Asor JE, Emmy-Egbe IO, et al. Epidemiology of tree-hole breeding mosquitoes in the tropical rainforest of Imo State, South-East Nigeria. Pediatr Dev Pathol. 1998;1: 200–209. doi:10.1007/s100249900027

24. Guindo-Coulibaly N, Adja AM, Koudou BG, Konan YL, Diallo M, Koné AB, et al. Distribution and seasonal variation of *Aedes aegypti* in the health district of Abidjan (Côte d'Ivoire). Eur J Sci Res. 2010;40: 522–530.
25. Bang YH, Knudsen AB, Onwubiko AO, Bown DN. Seasonal survival of *Aedes africanus* (Diptera: Culicidae) in Nigeria. J Med Entomol. 1983;20: 128–133. doi:10.1093/jmedent/20.2.128
26. McCrae AWR, Kirya BG. Yellow fever and Zika virus epizootics and enzootics in Uganda. Trans R Soc Trop Med Hyg. 1982;76: 552–562.
27. Cordellier R, Botjchité B, Roche J-C, Monteny N. Circulation selvatique du virus dengue 2 en 1980, dans les savanes sub-soudaniennes de Côte d'Ivoire. Ent med Parasitol. 1983;21: 165–179. Available: https://core.ac.uk/download/pdf/39874081.pdf
28. Darsie F., Richard Pradhan SP, Vaidya RG. Notes on the Mosquitoes of Nepal I . New Country Records and Revised *Aedes* Keys (Diptera, Culicidae). Seta. 1991;23: 39–45.
29. Harinasuta C, Sucharit S, Deesin T, Surathin K, Vutikes S. Bancroftian filariasis in Thailand, a new endemic area. J Trop Med Pub Hlth Seas. 1970;1: 233–245.
30. Young KI, Mundis S, Widen SG, Wood TG, Tesh RB, Cardosa J, et al. Abundance and distribution of sylvatic dengue virus vectors in three different land cover types in Sarawak, Malaysian Borneo. Parasites and Vectors. 2017;10: 1–14. doi:10.1186/s13071-017-2341-z
31. Wiwatanaratanabutr I. Geographic distribution of wolbachial infections in mosquitoes from Thailand. J Invertebr Pathol. 2013;114: 337–340. doi:10.1016/j.jip.2013.04.011
32. Santiago ATA, Claveria FG. Medically important mosquitoes (Diptera: Culicidae) identified in rural Barangay Binubusan, Lian, Batangas Province, Philippines. Philipp J Sci. 2012;141: 103–109.
33. Rogozi E, Ahmad RB, Ismail Z. Distribution and species composition of mosquitoes in three malay recreational parks. Burazeri G, Kakarriqi E, editors. Albanian Med J. 2012;4: 42–55. Available: http://www.ishp.gov.al/wp-content/uploads/2012/12/revista_nr_4_2012.pdf#page=30
34. Parker OS, Chaney AH. *Liomys irroratus* (Rodentia: Heteromyidae), a new host for *Cuterebra fontinella* (Diptera: Cuterebridae). J Med Entomol. 1979;15: 573–576. doi:10.1093/jmedent/15.5-6.573
35. Harrison BA, Rattanarithikul R, Peyton EL, Mongkolpanya K. Taxonomic changes, revised occurrence records and notes on the Culicidae of Thailand and neighboring countries. Mosq Syst. 1990;22: 196–227. Available: http://oai.dtic.mil/oai/oai?verb=getRecord&metadataPrefix=html&identifier=ADA512869
36. Chen CD, Lee HL, Stella-Wong SP, Lau KW, Sofian-Azirun M. Container survey of mosquito breeding sites in a university campus in Kuala Lumpur, Malaysia. Dengue Bull. 2009;33: 187–193.
37. Azmi NNM, Saad AR. Spatial distribution and habitat characterization of aedes mosquito larvae in Dengue hotspot areas on Penang Island. The 3rd International PSU-UNS Conferences on Bioscience. 2010. pp. 134–136.
38. Gould DJ, Bailey CL, Vongpradist S. Implication of forest mosquitoes in the transmission of *Wuchereria Bancrofti* in Thailand. Mosq News. 1982;42: 560–563.
39. Darsie F. Richard J, Gregory WC, Shreedhar PP. Notes on the mosquitoes of Nepal: III. Additional New Records in 1992 (Diptera: Culicidae). Mosq Syst. 1993;25: 186–191.
40. Richard, F. Dardsie J, Pradhan PP, Riddhi VG. Notes on the Mosquitoes of Nepal: II. New Species records from 1991 collections. Mosq Syst. 1992;24: 23–28.
41. Sucharit S, Rongsriyam Y, Deesin V, Komalamisra N, Apiwathnasorn C, Surathint K. Biology of Dengue Vectors and Their Control in Thailand. Trop Med. 1993;35: 253–257.
42. Ramírez JE, Yanoviak SP, Lounibos LP, Weaver SC. Distribución vertical de *Haemagogus janthinomys* (Dyar) (DIPTERA: CULICIDAE) en bosques de la Amazonía peruana. Rev Peru Med Exp Salud Publica. 2007;24:40–5.
43. Alencar J, De Mello VS, Serra-Freire NM, Silva JDS, Morone F, Guimarães AÉ. Evaluation of mosquito (Diptera: Culicidae) species richness using two sampling methods in the hydroelectric reservoir of Simplício, Minas Gerais, Brazil. Zoolog Sci. 2012;29:218–22.
44. Alencar J, de Mello CF, Gil-Santana HR, Guimarães AÉ, de Almeida SAS, Gleiser RM. Vertical oviposition activity of mosquitoes in the Atlantic Forest of Brazil with emphasis on the sylvan vector, *Haemagogus leucocelaenus* (Diptera: Culicidae). J Vector Ecol. 2016;41:18–26.
45. Ramírez JE, Yanoviak SP, Lounibos LP, Weaver SC. Distribución vertical de *Haemagogus janthinomys* (Dyar) (DIPTERA: CULICIDAE) en bosques de la Amazonía peruana. Rev Peru Med Exp Salud Publica. 2007;24:40–5.
46. de Almeida MAB, dos Santos E, Cardoso J da C, da Silva LG, Rabelo RM, Bicca-Marques JC. Predicting Yellow Fever Through Species Distribution Modeling of Virus, Vector, and Monkeys. Ecohealth. 2019;16:95–108.

**Table S2.** Variables of the logit functions of the baseline models in Aliaga-Samanez *et al*. 2021, 2022. Logit functions are linear combinations (*y*) of predictor variables that form part of the favourability (*F*) function: *F* = exp(*y*) / [(*n1*/*n0*) + exp(*y*)]. In the case of *Ae. aegypti* and *Ae. albopictus*, the predictor variable "*Y-20$^{th}$ century*" is the logit equation of the 20$^{th}$ century model (see Aliaga-Samanez et al. 2021, 2022). Description and sources of predictor variables can be found in table S4. B: variable coefficient; SE: standard error; W: Wald parameter; DF: degrees of freedom; S: statistical significance.

| ***Aedes aegypti*** | | | | | |
|---|---|---|---|---|---|
| **Variables** | B | SE | W | DF | S |
| ***Y-20$^{th}$ century*** | 0.096 | 0.048 | 3.949 | 1 | 0.047 |
| ***Bio12*** | $-0.207x10^{-3}$ | $0.65x10^{-4}$ | 10.235 | 1 | 0.001 |
| ***Bio6*** | 0.010 | 0.001 | 137.132 | 1 | $0.113x10^{-30}$ |
| ***Class 110*** | -2.715 | 0.922 | 8.678 | 1 | 0.003 |
| ***Class 150*** | -2.273 | 0.994 | 5.228 | 1 | 0.022 |
| ***Class 160*** | -2.690 | 0.886 | 9.211 | 1 | 0.002 |
| ***Class 200*** | -1.483 | 0.389 | 14.538 | 1 | $0.137x10^{-3}$ |
| ***Class 50*** | -1.271 | 0.404 | 9.899 | 1 | 0.002 |
| ***Class 70*** | 1.336 | 0.573 | 5.442 | 1 | 0.020 |
| ***Dist_pop*** | $-0.1x10^{-4}$ | $0.315x10^{-5}$ | 10.161 | 1 | 0.001 |
| ***Equi_irrig*** | 0.015 | 0.005 | 9.578 | 1 | 0.002 |
| ***Pop_den*** | 0.001 | $0.158x10^{-3}$ | 69.113 | 1 | $0.929x10^{-16}$ |
| ***Poultry*** | $0.72x10^{-4}$ | $0.627x10^{-4}$ | 8.694 | 1 | 0.003 |
| ***Sheep*** | -0.008 | 0.003 | 6.286 | 1 | 0.012 |
| ***Slope*** | 0.111 | 0.026 | 18.056 | 1 | $0.214x10^{-4}$ |
| ***FAfrica*** | 1.071 | 0.241 | 19.747 | 1 | $0.884x10^{-5}$ |
| ***FAsia*** | 1.016 | 0.209 | 23.579 | 1 | $0.112x10^{-5}$ |
| ***FAmericanorth*** | 1.838 | 0.251 | 53.673 | 1 | $0.237x10^{-12}$ |
| ***FAmericasouth*** | 6.222 | 0.223 | 780.911 | 1 | $0.763x10^{-171}$ |
| ***FEurope*** | 6.601 | 2.613 | 6.380 | 1 | 0.012 |
| ***Constant*** | -4.034 | 0.248 | 263.853 | 1 | $0.248x10^{-58}$ |
| ***Aedes albopictus*** | | | | | |
| **Variables** | B | SE | W | DF | S |
| ***Y-20$^{th}$ century*** | 0.089 | 0.029 | 9.124 | 1 | 0.003 |
| ***Bio15*** | 0.009 | 0.002 | 20.732 | 1 | $0.528x10^{-5}$ |
| ***Bio12*** | $0.341x10^{-3}$ | $0.909x10^{-4}$ | 14.081 | 1 | $0.175x10^{-3}$ |
| ***Class 130*** | -1.526 | 0.413 | 13.672 | 1 | $0.218x10^{-3}$ |
| ***Class 200*** | -7.853 | 2.891 | 7.382 | 1 | 0.007 |
| ***Class 40*** | 1.118 | 0.275 | 16.548 | 1 | $0.474x10^{-4}$ |
| ***Dist_rail*** | $-0.108x10^{-5}$ | $0.465x10^{-6}$ | 5.387 | 1 | $0.203x10^{-1}$ |
| ***Dist_road*** | $-0.406x10^{-5}$ | $0.169x10^{-5}$ | 5.743 | 1 | $0.166x10^{-1}$ |
| ***Dist_pop*** | $-0.135x10^{-4}$ | $0.359x10^{-5}$ | 14.144 | 1 | $0.169x10^{-3}$ |
| ***Pop_den*** | 0.002 | $0.163x10^{-3}$ | 107.589 | 1 | $0.331x10^{-24}$ |
| ***Pigs*** | 0.002 | 0.001 | 4.435 | 1 | 0.035 |

| | | | | | |
|---|---|---|---|---|---|
| ***FAfrica*** | 3.219 | 0.281 | 131.037 | 1 | 0.243x10$^{-29}$ |
| ***FAsia*** | 2.350 | 0.325 | 52.336 | 1 | 0.468x10-$^{12}$ |
| ***FOceania*** | 3.215 | 0.569 | 31.875 | 1 | 0.164x10$^{-7}$ |
| ***FAmericanorth*** | 4.532 | 0.360 | 158.799 | 1 | 0.207x10$^{-35}$ |
| ***FAmericasouth*** | 7.891 | 0.251 | 985.359 | 1 | 0.273x10$^{-215}$ |
| ***FEurope*** | 6.099 | 0.291 | 440.676 | 1 | 0.771x10$^{-97}$ |
| ***Constant*** | -5.497 | 0.339 | 263.690 | 1 | 0.268x10$^{-58}$ |
| ***Aedes africanus*** | | | | | |
| Variables | B | SE | W | DF | S |
| ***Bio15*** | 0.024 | 0.006 | 14.666 | 1 | 0.128X10$^{-3}$ |
| ***Bio12*** | 0.001 | 0.249X10$^{-3}$ | 8.88 | 1 | 0.003 |
| ***Bio6*** | 0.024 | 0.006 | 13.762 | 1 | 0.207X10$^{-3}$ |
| ***Class 130*** | 2.731 | 1.19 | 5.266 | 1 | 0.022 |
| ***Class 30*** | 5.707 | 0.898 | 40.344 | 1 | 0.213X10$^{-9}$ |
| ***Class 60*** | 6.077 | 1.053 | 33.303 | 1 | 0.789X10$^{-8}$ |
| ***Pop_den*** | 0.001 | 0.337X10$^{-3}$ | 3.035 | 1 | 0.081 |
| ***Goats*** | 0.012 | 0.004 | 8.971 | 1 | 0.003 |
| ***Sheep*** | 0.016 | 0.004 | 13.398 | 1 | 0.252X10$^{-3}$ |
| ***Constant*** | -15.508 | 1.751 | 78.463 | 1 | 0.815X10$^{-18}$ |
| ***Aedes luteocephalus*** | | | | | |
| Variables | B | SE | W | DF | S |
| ***Bio6*** | 0.035 | 0.010 | 12.199 | 1 | 0.478X10$^{-3}$ |
| ***Bio5*** | 0.031 | 0.009 | 12.264 | 1 | 0.462X10$^{-3}$ |
| ***Class 130*** | 3.806 | 1.355 | 7.889 | 1 | 0.005 |
| ***Class 30*** | 6.752 | 1.236 | 29.838 | 1 | 0.470X10$^{-7}$ |
| ***Class 60*** | 6.486 | 1.299 | 24.931 | 1 | 0.594X10$^{-6}$ |
| ***Goats*** | 0.017 | 0.004 | 20.117 | 1 | 0.7X10$^{-5}$ |
| ***Pop_den*** | 0.001 | 0.362X10$^{-3}$ | 7.270 | 1 | 0.007 |
| ***Constant*** | -25.799 | 4.553 | 32.113 | 1 | 0.145X10$^{-7}$ |
| ***Aedes niveus*** | | | | | |
| Variables | B | SE | W | DF | S |
| ***Bio12*** | 0.001 | 0.259X10$^{-3}$ | 7.818 | 1 | 0.005 |
| ***Bio6*** | 0.017 | 0.004 | 14.87 | 1 | 0.115X10$^{-3}$ |
| ***Class 11-14*** | 4.736 | 1.483 | 10.204 | 1 | 0.001 |
| ***Equi_irrig*** | 0.054 | 0.019 | 8.377 | 1 | 0.004 |
| ***Slope*** | 0.687 | 0.128 | 28.791 | 1 | 0.806X10$^{-7}$ |
| ***Constant*** | -13.729 | 1.485 | 85.494 | 1 | 0.232X10$^{-19}$ |
| ***Aedes vittatus*** | | | | | |
| Variables | B | SE | W | DF | S |
| ***Bio6*** | 0.012 | 0.002 | 28.833 | 1 | 0.789X10$^{-7}$ |
| ***Buffaloes*** | 0.021 | 0.006 | 12.646 | 1 | 0.376X10$^{-3}$ |
| ***Class 130*** | 2.513 | 0.754 | 11.117 | 1 | 0.001 |
| ***Class 30*** | 2.428 | 0.815 | 8.864 | 1 | 0.003 |

| | | | | | |
|---|---|---|---|---|---|
| ***Dist_rail*** | $-0.13X10^{-4}$ | $0.4X10^{-5}$ | 12.845 | 1 | $0.338X10^{-3}$ |
| ***Goats*** | 0.007 | 0.003 | 7.229 | 1 | 0.007 |
| ***Sheep*** | 0.008 | 0.003 | 7.472 | 1 | 0.006 |
| ***Constant*** | -7.24 | 0.465 | 242.182 | 1 | $0.131X10^{-53}$ |
| ***Sabethes chloropterus*** | | | | | |
| **Variables** | B | SE | W | DF | S |
| ***Bio6*** | 0,012 | 0,004 | 7,349 | 1 | 0,007 |
| ***Class30*** | 2,718 | 1,228 | 4,896 | 1 | 0,027 |
| ***Class40*** | 1,4 | 0,762 | 3,373 | 1 | 0,066 |
| ***Constant*** | -8,99 | 0,773 | 135,076 | 1 | 0 |
| ***Haemagogus janthinomys*** | | | | | |
| **Variables** | B | SE | W | DF | S |
| ***Bio7*** | 0,002 | 0,006 | 0,194 | 1 | 0,659 |
| ***Class40*** | 1,624 | 0,842 | 3,717 | 1 | 0,054 |
| ***Cattle*** | 0,008 | 0,009 | 0,921 | 1 | 0,337 |
| ***FAmerica*** | 7,635 | 1,488 | 26,33 | 1 | 0 |
| ***Constant*** | -11,285 | 1,771 | 40,603 | 1 | 0 |
| ***Haemagogus leucocelaenus*** | | | | | |
| **Variables** | **B** | **SE** | **W** | **DF** | **S** |
| ***Bio15*** | -0,053 | 0,011 | 22,532 | 1 | 0 |
| ***Bio5*** | 0,019 | 0,008 | 6,103 | 1 | 0,013 |
| ***Class30*** | 2,947 | 1,562 | 3,56 | 1 | 0,059 |
| ***Class20*** | 8,347 | 1,383 | 36,418 | 1 | 0 |
| ***Class40*** | 6,314 | 1,015 | 38,704 | 1 | 0 |
| ***Dist_rail*** | $-0.1X10^{-4}$ | 0 | 8,153 | 1 | 0,004 |
| ***Cattle*** | 0,017 | 0,004 | 21,718 | 1 | 0 |
| ***Constant*** | -12,496 | 2,205 | 32,131 | 1 | 0 |

**Table S3.** Variables in the logit equations of the vector models (*Aedes aegypti* and *Aedes albopictus*) of the 20th century (i.e., linear combinations of predictor variables that form part of the logistic-regression equations). Description and sources of predictor variables can be found in table S4. B: variable coefficient; SE: standard error; W: Wald parameter; DF: degrees of freedom; S: statistical significance.

| 20th-century models | | | | | |
|---|---|---|---|---|---|
| ***Aedes aegypti*** | | | | | |
| **Variable** | **B** | **SE** | **W** | **DF** | **S** |
| ***Bio12*** | $0.215 \times 10^{-3}$ | $0.723 \times 10^{-4}$ | 8.854 | 1 | 0.003 |
| ***Bio5*** | $0.568 \times 10^{-2}$ | 0.002 | 13.581 | 1 | $0.228 \times 10^{-3}$ |
| ***Dist_pop*** | $-0.483 \times 10^{-4}$ | $0.476 \times 10^{-5}$ | 103.018 | 1 | $0.332 \times 10^{-23}$ |
| ***Elev*** | $-0.342 \times 10^{-3}$ | $0.122 \times 10^{-3}$ | 7.828 | 1 | 0.005 |
| ***FAmericanorth*** | 5.103 | 0.233 | 478.237 | 1 | $0.517 \times 10^{-105}$ |
| ***FAmericasouth*** | 4.765 | 0.293 | 264.171 | 1 | $0.211 \times 10^{-58}$ |
| ***FAfrica*** | 4.230 | 0.284 | 222.491 | 1 | $0.259 \times 10^{-49}$ |
| ***FAsia*** | 3.953 | 0.262 | 228.428 | 1 | $0.131 \times 10^{-50}$ |
| ***FOceania*** | 5.543 | 0.310 | 318.722 | 1 | $0.275 \times 10^{-70}$ |
| ***Constant*** | -6.277 | 0.539 | 135.832 | 1 | $0.217 \times 10^{-30}$ |
| ***Aedes albopictus*** | | | | | |
| **Variable** | **B** | **SE** | **W** | **DF** | **S** |
| ***Bio12*** | 0.001 | $0.117 \times 10^{-3}$ | 20.335 | 1 | $0.650 \times 10^{-5}$ |
| ***Bio7*** | 0.003 | 0.001 | 8.958 | 1 | $0.276 \times 10^{-2}$ |
| ***Dist_pop*** | $-0.334 \times 10^{-4}$ | $0.653 \times 10^{-5}$ | 26.154 | 1 | $0.315 \times 10^{-6}$ |
| ***Elev*** | -0.001 | $0.220 \times 10^{-3}$ | 13.598 | 1 | $0.226 \times 10^{-3}$ |
| ***TempCF*** | 1.242 | 0.378 | 10.801 | 1 | 0.001 |
| ***FAmericanorth*** | 8.533 | 0.371 | 529.728 | 1 | $0.324 \times 10^{-116}$ |
| ***FAmericasouth*** | 6.204 | 0.451 | 189.334 | 1 | $0.444 \times 10^{-42}$ |
| ***FEurope*** | 6.186 | 0.442 | 195.751 | 1 | $0.176 \times 10^{-43}$ |
| ***FAfrica*** | 5.366 | 0.533 | 101.278 | 1 | $0.799 \times 10^{-23}$ |
| ***FAsia*** | 5.429 | 0.458 | 140.438 | 1 | $0.213 \times 10^{-31}$ |
| ***FOceania*** | 6.235 | 0.538 | 134.205 | 1 | $0.493 \times 10^{-30}$ |
| ***Constant*** | -7.590 | 0.479 | 250.819 | 1 | $0.172 \times 10^{-56}$ |

**Table S4.** Predictor variables considered for baseline model training in Aliaga-Samanez *et al*. 2021, 2022.

| Factor | Code | Variable | Source |
|---|---|---|---|
| Climate | *Bio1* | Annual Mean Temperature | Chelsa (http://chelsa-climate.org) |
| | *Bio5* | Max Temperature of Warmest Month | |
| | *Bio6* | Min Temperature of Coldest Month | |
| | *Bio7* | Temperature Annual Range (Bio5-Bio6) | |
| | *Bio12* | Annual Precipitation | |
| | *Bio15* | Precipitation Seasonality (Coefficient of Variation) | |
| Human Concentration | *Pop_den* | Population density | LandScan™ 2008 High Resolution Global Population (https://landscan.ornl.gov/) |
| | *Dist_pop* | Distance to populated places | Administrative Centres & Populated Places shapefile at the Relational World Database II (RWDB2) updated in 2000 (http://www.fao.org/geonetwork) |
| Infrastructures | *Dist_road* | Distance to roads | Vector Map Level 0 at the Digital Chart of the World (DCW,http://worldmap.harvard.edu), updated in 2002 |
| | *Dist_rail* | Distance to rail-roads | |
| Livestock | *Buffaloes* | Density of buffaloes | FAO 2010(http://www.fao.org/livestock-systems/en/) |
| | *Poultry* | Density of poultry | |
| | *Goats* | Density of small ruminants (goats) | |
| | *Pigs* | Density of pigs | |
| | *Sheep* | Density of small ruminants (sheep) | |
| | *Cattle* | Density of cattle | |
| Topography | *Slope* | slope | From GTOPO30 (US Geological Survey 1996), using ArcGIS Desktop 10.3. |
| | *Elev* | Elevation | GTOPO30 (US Geological Survey 1996). |
| Ecoregions | *TempCF* | Temperate Coniferous Forests | Terrestrial Ecoregions of the World: A New Map of Life on Earth: A new global map of terrestrial ecoregions provides an innovative tool for conserving biodiversity [47] |
| Agriculture | *Class 11-14* | Croplands | GlobCover (GC) Land Cover version 2.3 database for 2009 [48] |
| | *Class 20* | Mosaic Cropland (50-70%) / Vegetation (grassland, shrubland, forest) (20-50%) | |
| | *Class 30* | Mosaic Vegetation (grassland, shrubland, forest) (50-70%) / Cropland (20-50%) | |
| | *Equi_irrig* | Percentage of area equipped for irrigation | Global Map of Irrigation Areas (version 4.0.1) around the year 2000 (http://www.fao.org/nr/water) |
| Ecosystem Types | *Class 40* | Closed to open (>15%) broadleaved evergreen | |

| | | | |
|---|---|---|---|
| | | and/or semi-deciduous forest (>5m) | GlobCover (GC) Land Cover version 2.3 database for 2009 [48] |
| | *Class 50* | Closed (>40%) broadleaved deciduous forest (>5m) | |
| | *Class 60* | Open (15-40%) broadleaved deciduous forest (>5m) | |
| | *Class 70* | Closed (>40%) needleleaved evergreen forest (>5m) | |
| | *Class 110* | Mosaic Forest/Shrubland (50-70%) / Grassland (20-50%) | |
| | *Class 130* | Closed to open (>15%) shrubland (<5m) | |
| | *Class 150* | Sparse (>15%) vegetation (woody vegetation, shrubs, grassland) | |
| | *Class 160* | Closed (>40%) broadleaved semi-deciduous and/or evergreen forest regularly flooded - Saline water | |
| | *Class 200* | Bare areas | |
| Logit equation | *Y-20$^{th}$century* | 20$^{th}$-century-model logit equation | Linear combinations of predictor variables that form part of the logistic-regression equations |
| Spatial descriptors | *Fx* | Spatial trend, where "x" represents a continent | Linear combination of spatial variables derived from continental-scale trend surface analyses [49] |


**References:**

47. Olson DM, Dinerstein E, Wikramanayake E, Burgess ND, Powell G, Underwood E, et al. Terrestrial Ecoregions of the World: A New Map of Life on Earth: A new global map of terrestrial ecoregions provides an innovative tool for conserving biodiversity. Bioscience. 2001; 51: 933–938.
48. Bontemps S. GLOBCOVER 2009 Products Description and Validation Report. 2011.
49. Legendre P. Spatial autocorrelation: Trouble or New Paradigm? Ecology. 1993;74: 1659–1673

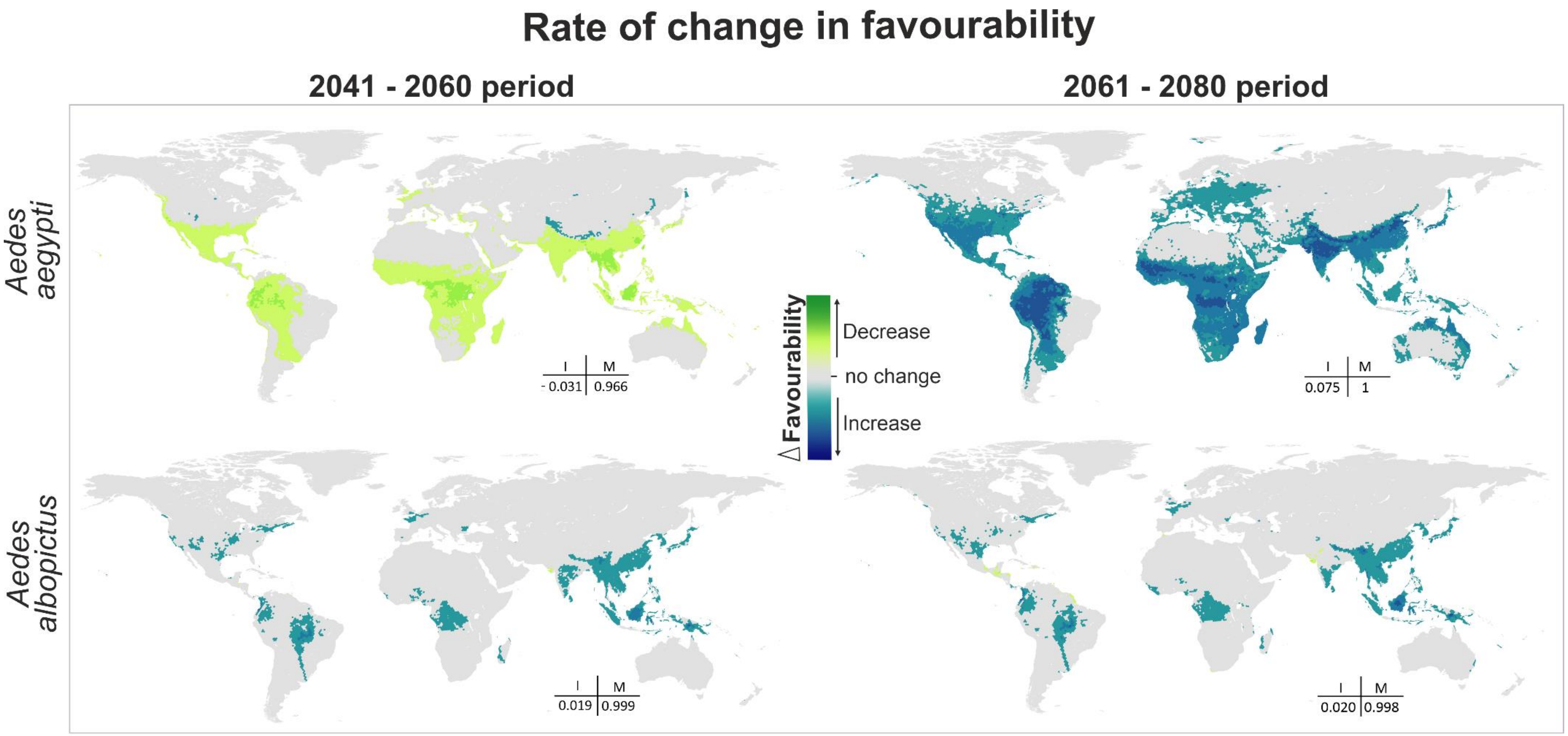


**Fig. S1. Areas where favourability increases and decreases in the future relative to the present.** Difference between the future projection and the current model. I: increment rate; B: maintenance rate (Romero et al. Journal of Biogeography 41.1 (2014): 111-121). Positive values of I indicate a net increase in favourability, that is, a gain in favourable areas, whereas negative values of I mean a net loss of favourable areas. M indicates the degree to which the favourable areas in the current model overlap with the favourable forecasted area.

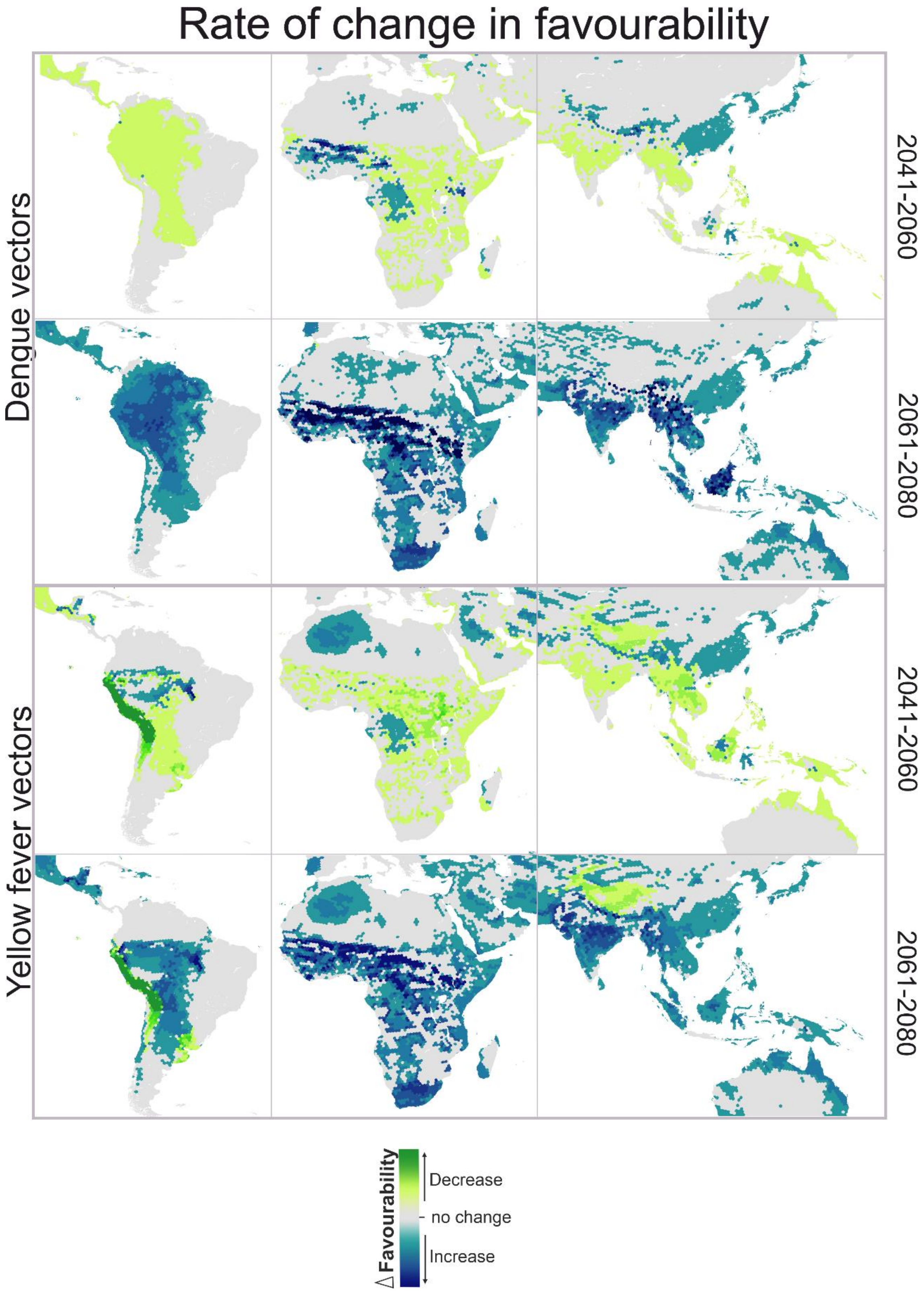


**Fig. S2. Rate of change in favourability of dengue and yellow fever vectors models.** Zoomed details of areas where favourability increases and decreases in the future relative (2041-2060 and 2061 – 2080) to the present**.**
